# Islands in the desert: Environmental distribution modelling of endemic flora reveals the extent of Pleistocene tropical relict vegetation in southern Arabia

**DOI:** 10.1101/563650

**Authors:** James S. Borrell, Ghudaina Al Issaey, Darach A. Lupton, Thomas Starnes, Abdulrahman Al Hinai, Saif Al Hatmi, Rebecca A. Senior, Tim Wilkinson, Jo L.H. Milborrow, Andrew Stokes-Rees, Annette Patzelt

## Abstract

**Background and Aims:** Southern Arabia is a global biodiversity hotspot with a high proportion of endemic desert-adapted plants. Here we examine evidence for a Pleistocene climate refugium in the southern Central Desert of Oman, and its role in driving biogeographical patterns of endemism.

**Methods:** Distribution data for seven narrow-range endemic plants were collected systematically across 195 quadrats, together with incidental and historic records. Important environmental variables relevant to arid coastal areas, including night time fog and cloud cover were developed for the study area. Environmental niche models were built and tuned for each species and spatial overlap examined.

**Key Results:** A region of the Jiddat Al Arkad reported independent high model suitability for all species. Examination of environmental data across southern Oman indicates that the Jiddat Al Arkad displays a regionally unique climate with higher intra-annual stability, due in part to the influence of the southern monsoon. Despite this, relative importance of environmental variables was highly differentiated among species, suggesting characteristic variables such as coastal fog are not major cross-species predictors at this scale.

**Conclusions:** The co-occurrence of a high number of endemic study species within a narrow monsoon-influenced region is indicative of a refugium with low climate change velocity. Combined with climate analysis, our findings provide strong evidence for a southern Arabian Pleistocene refugium in the Oman’s Central Desert. We suggest this refugium has acted as an isolated temperate and mesic island in the desert, resulting in the evolution of these narrow-range endemic flora. Based on the composition of species, this system may represent the northernmost remnant of a continuous belt of mesic vegetation formerly ranging from Africa to Asia, with close links to the flora of East Africa. This has significant implications for future conservation of endemic plants in an arid biodiversity hotspot.

## INTRODUCTION

Southern Arabia is part of the Horn of Africa global biodiversity hotspot, and is one of only two hotspots that are entirely arid (Mittermeier *et al.* 2005; Mallon 2013). The flora of southern Arabia arises from the relatively late separation of Arabia from Africa and Asia around 25 million years before present (Raven and Axelrod 1974; Delany 1989). During the Miocene Arabia supported palaeo-tropical vegetation with swamps and open savannah grassland (Whybrow and Mcclure 1981). This was progressively replaced by more drought-adapted vegetation through the Pliocene, with mesic elements of the flora persisting only in climatically favourable refugia (Kürschner 1998; Jolly *et al.* 2009). The environment of southern Arabia subsequently oscillated between climatic extremes throughout the Quaternary period (Fleitmann and Matter 2009; Parker 2010; Jennings *et al.* 2015). These oscillations, combined with the relative stability of localized climatic refugia may have contributed to the high degree of species endemism (Patzelt, 2015; Sandel et al., 2011).

The biogeographic consequences of contraction and expansion from glacial refugia have been well described in the temperate zones of Europe and North America (Bennett *et al.* 1991; Comes and Kadereit 1998; Birks and Willis 2008; Keppel *et al.* 2012; Wang *et al.* 2014). By comparison, these processes are poorly known in the arid environments of the Arabian Peninsula (Ghazanfar, 1998; Meister, Hubaishan, Kilian, & Oberprieler, 2006; Patzelt, 2015).

Therefore establishing the spatiotemporal distribution of past climate refugia in southern Arabia is likely to have important implications for future conservation planning (Al-Abbasi *et al.* 2010), building evolutionary resilience under climate change (Sgrò *et al.* 2011; Keppel *et al.* 2012) and even interpreting the history of hominid dispersal out of Africa (Jennings *et al.* 2015; Gandini *et al.* 2016).

A key center for plant endemism in southern Arabia is Oman’s Central Desert (Ghazanfar, 2004; Miller & Nyberg, 1990; Patzelt, 2014; White & Léonard, 1990). The Central Desert is a hyper-arid region, characterized by scarce precipitation often less than 100 mm/pa with high inter-annual variability and temperatures ranging from 6°C to more than 50°C (Stanley Price *et al.* 1988; Fisher and Membery 1998; Almazroui *et al.* 2013). Provisionally divided into ‘northern’ and ‘southern’ systems, the Central Desert has relatively low species diversity, but the highest proportion of range restricted endemic and regionally endemic plants in Oman (Patzelt, 2015). This represents an ideal study system in which to test for evidence of climatic refugia and their influence on the floral biogeography of southern Arabia.

Despite significant progress in documenting the flora of Oman (Brinkmann et al., 2011; Ghazanfar, 1998, 2004; Ghazanfar & Fisher, 2013; Miller & Cope, 1996; Patzelt, 2009, 2014) high resolution plant diversity and distribution data are limited or lacking for many areas, hindering our ability to test these biogeographic hypotheses and identify putative refugia. To address this knowledge gap, here we report results of a systematic botanic survey of the southern Central Desert. We focus on seven high priority narrow range endemic desert plants (Table 1; Figure 1), restricted to the coastal belt and the adjacent escarpment and identified through development of the Oman Red Data Book (Patzelt, 2014).

**Table 1.**
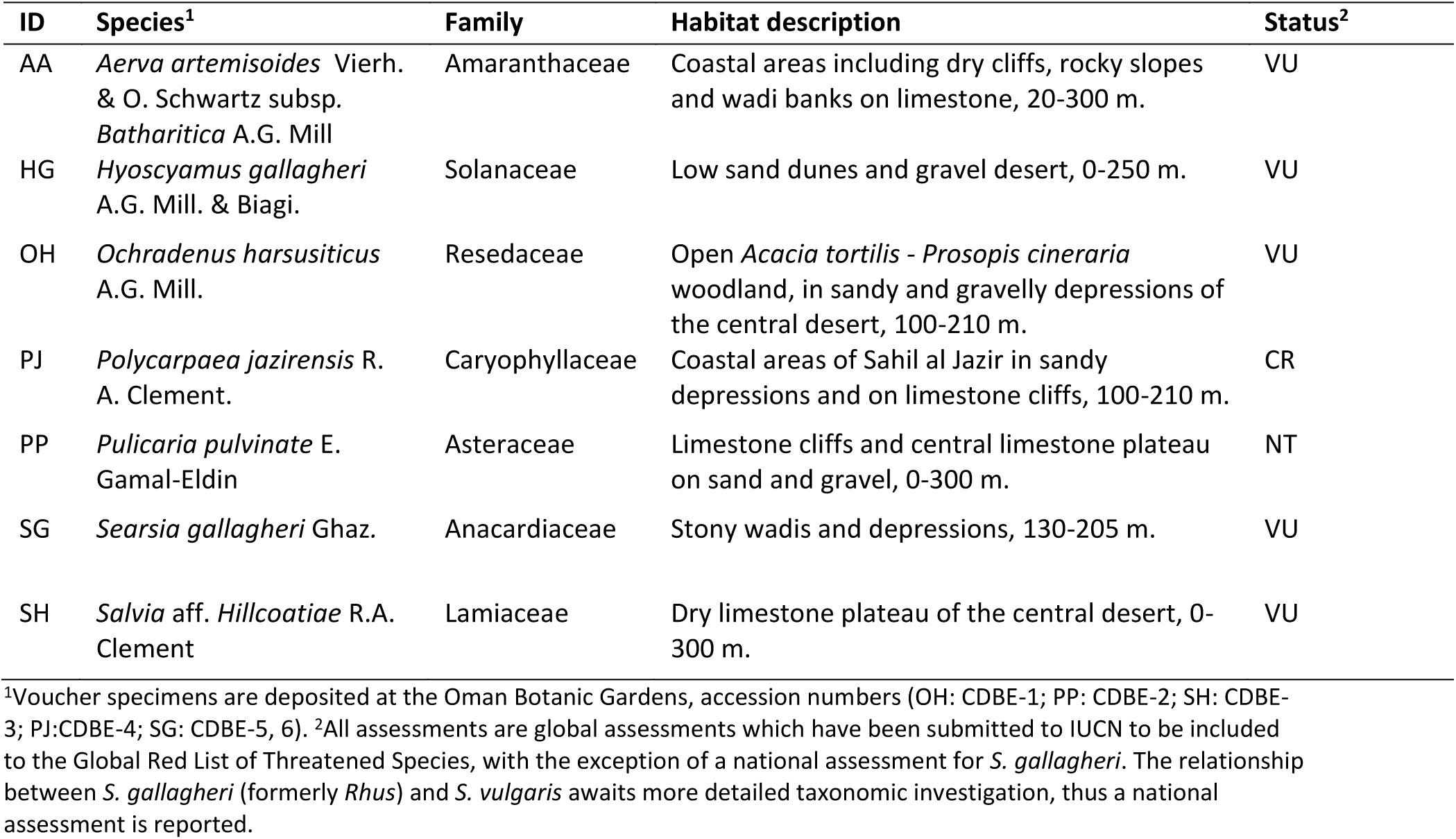
Summary of the seven study taxa, including Red List status from Patzelt (2014).

**Figure 1.**
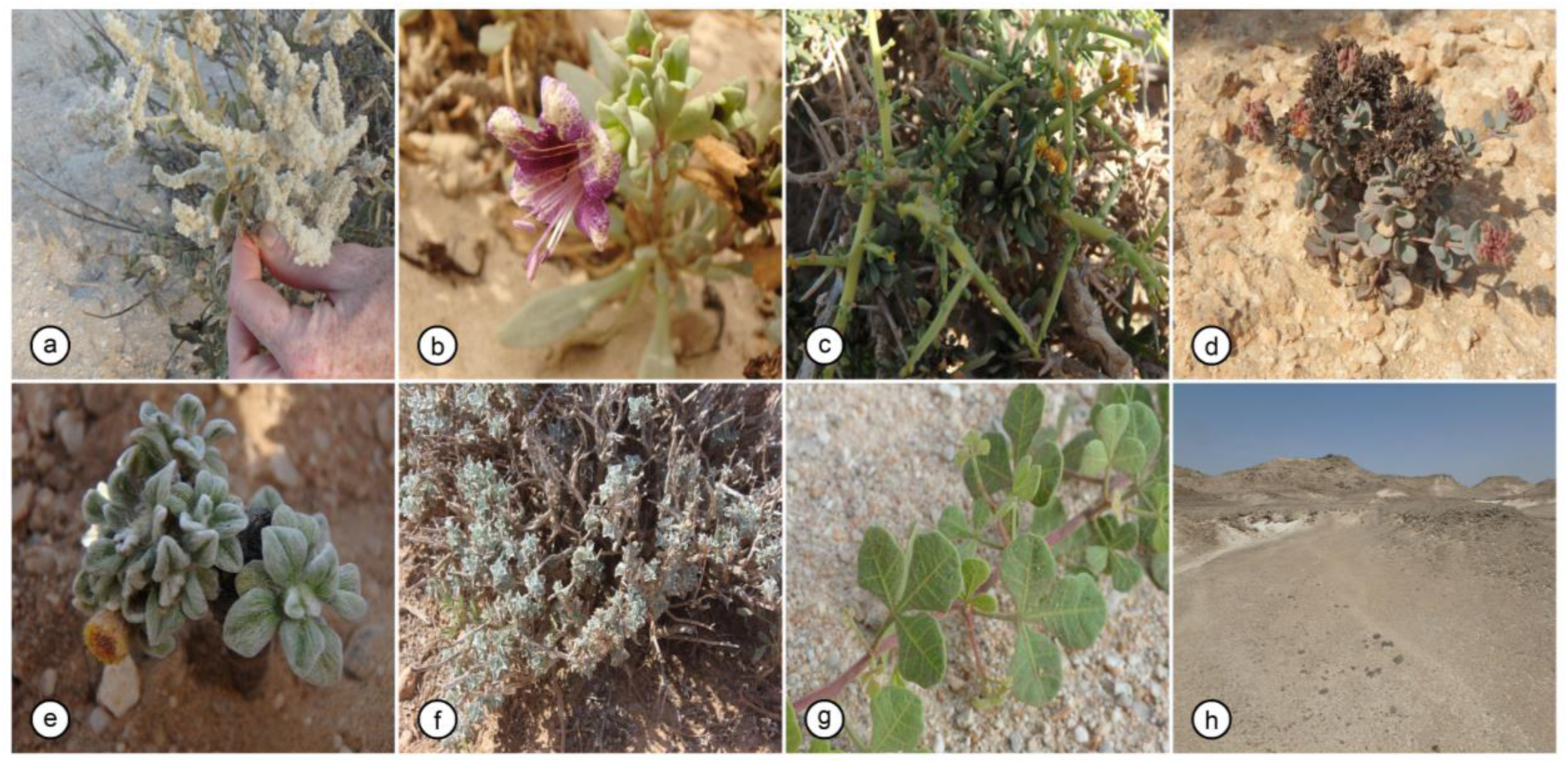
Images of the endemic Central Desert plant species included in this study. a) *Aerva artemisoides subsp. batharitica*; b) *Hyoscyamus gallagheri*; c) *Ochradenus harsusiticus*; d) *Polycarpaea jazirensis*; e) *Pulicaria pulvinata*; f) *Salvia aff. hillcoatiae*; g) *Searsia gallagheri*; h) A typical Central Desert landscape.

To enable a transition from survey data to predictive distribution maps we use an environmental niche modelling (ENM) approach. ENMs are a suite of methods used to establish the relationship between a species and a set of environmental variables (Elith and Leathwick, 2009; Peterson, Papeş and Soberón, 2015). In principle, ENMs evaluate the environmental conditions in grid cells known to be occupied by a species and identify additional cells that represent similar environmental conditions (Merow, Smith and Silander, 2013). The species’ niche can then be projected across the study area to predict its spatial distribution and identify environmental variables that contribute significantly to model performance (Searcy and Shaffer 2016). In this study we benefit from both presence and absence survey data and a stratified survey design, which negates many of the biases common in environmental niche models (Jiménez-Valverde *et al.* 2008; Warren and Seifert 2011).

To distinguish amongst alternative environmental drivers for local endemism, selection of appropriate environmental modelling variables in arid environments is important (Dilts *et al.* 2015; Title and Bemmels 2018). In other coastal desert systems, endemic plant distributions are strongly influenced by the presence of cloud shade and fog, which causes condensation on leaves and stems that trickles down to root systems (Cereceda *et al.* 2008; Fischer *et al.* 2009). Given the extreme temperatures and low precipitation, the presence of coastal fog and the cooling effect of the prevailing wind have been hypothesised to be a key driver of Central Desert flora distributions (Miller, 1994; Patzelt, 2015; Price et al., 1988). The presence of coastal fog is supported by data from the northern Central Desert in 1984, where water collected from fog collectors at Jiddat al Harasisi ranged from 0.08 L/m^2^ in January to 3.6 L/m^2^. A total of 93 nights with fog moisture were recorded across the year, with fog moisture at ground level coinciding with reduced night-time temperatures, increased humidity and a wind speed less than 15 Km/h (Price et al., 1988). In a subsequent study (Fisher and Membery 1998) a monthly maximum of 4.0 L/m^2^ during March and a minimum 2.5 L/m^2^ during January, May, June and December was recorded. To our knowledge, no empirical data is available on coastal fog from the southern central desert.

Validation of environmental variables across a large, arid and sparsely populated study area such as the Central Desert is also exacerbated by the paucity of weather stations (meaning that global climate models are highly interpolated) and the fact that where weather stations do exist, cloud cover and coastal fog are rarely recorded. To address these shortfalls, we make use of the newly available WorldClim2 dataset that incorporates high accuracy remotely sensed maximum and minimum land surface temperature (Fick and Hijmans 2017), together with remotely sensed cloud cover data (Wilson and Jetz 2016). We also incorporate remotely sensed fog and putative fog covariates, to explore whether these variables contribute significantly to Central Desert species distributions. Specifically, we derive night time fog intensity from MODIS data (MODIS Characterization Support Team (MCST) 2018) using the approach of Chaurasia et al. (2011), as well as topography (including elevation, slope, aspect and terrain roughness, as locally higher areas may catch more moisture (Schemenauer *et al.* 1987)), wind speed and night time land surface temperature (LST) (Wan, Hook, & Hulley, 2018). As an additional line of evidence in assessing the importance of coastal fog, we also survey physiological fog capture adaptations across our study species using the approach of Larraín-Barrios *et al.* (2018).

Here, building on novel systematic survey data from the Central Desert, we aim to address three main questions. First, we use newly available climate data to examine the influence of the southern monsoon and coastal fog influx on the Central Desert climate. Second, we model the distributions of seven high priority narrow-range endemics, and test the hypothesis that the same environmental variables are consistently important across taxa. Third, we consider evidence for past climatic refugia and their influence on the current floral biogeography of the Central Desert. We consider these data in the context of conserving rare desert endemics across southern Arabia, a global biodiversity hotspot.

## MATERIALS AND METHODS

### Study area

The southern Central Desert is dominated by the Jiddat Al Arkad, a meandering escarpment of 50 to 100 m dissected by extensive wadi systems, depressions and runnels which discharge into the Sahil Al Jazir coastal plain. Surface flows are only present following heavy rains. Geologically, the study area is dominated by Oligocene – Miocene white bioclastic limestone with coral debris flow deposits and laminated dolomitic limestone (Patel 1992). Soil is predominantly Calciorthids – gravelly sandy loam on alluvial fans and stream terraces and torriorthents (rock outcrops) weakly or undeveloped, low in organic matter and moderately calcareous (Dregne 1976).

The vegetation is classified into three units as per Patzelt, (2015): i) *Acacia tortilis* – *Prosopis cineraria* open woodland. Common grass and shrub species include the endemic shrubs, *Convolvulus oppositifolius* and *Ochradenus harsusiticus* and endemic grass *Stipagrostis sokotrana.* ii) Xeromorphic dwarf shrubland intermixed with grasses and annual species. The dwarf palm *Nannorrhops ritchieana* and *A. ehrenbergiana* are common in sandy depressions close to the coastal escarpments. iii) Xeromorphic dwarf shrubland with *Searsia gallagheri* and *O. harsusiticus*. In addition to flowering plants, several species of corticolous and saxicolous lichens and epilithic cyanobacteria occur here, of which most are restricted to the putatively fog-affected zones (Ghazanfar & Gallagher, 1998). The seven study species (Figure 1) are considered a part of, though not restricted to, the xeromorphic dwarf shrubland community. A description of their known habitat and conservation status is provided in Table 1.

### Field surveys and plant morphology

Fieldwork was conducted during the period 13^th^ – 24^rd^ January 2017, from Ras Madrakah, across the Sahil Al Jazer (coastal plain) to the southern extent of the Jiddat Al Arkad, as part of the Central Desert Botanic Expedition 2017. A stratified survey strategy was designed orientating ten 20 km transects at 315 degrees (NW) along a 270 km portion of coastline, at 30 km intervals. This approach was designed to cross multiple environmental gradients that frequently run perpendicular to the coastline. Stratified quadrat locations at 1 km intervals were plotted prior to field work and their coordinates uploaded to handheld GPS units (Garmin, Oregon). Due to the remote nature of the study area, with several deep wadis bisecting these transects and inhibiting access, some portions of these transects were not surveyed. When moving between transects we opportunistically sampled additional quadrats at 5 km intervals and recorded incidental observations of target species to maximise data collection. These additional quadrats were positioned via random number generation to determine distance and bearing from the vehicle. A significant portion of travel was away from roads, but where roads (mostly gravel tracks) were present, quadrat positioning began > 100 m from the road to mitigate disturbance bias in the vegetation recorded.

The following data were recorded for each quadrat: location, soil texture, soil pH, soil electrical conductivity (EC) (following the method of Zhang *et al.* 2005), total vegetation cover, maximum vegetation height, elevation, topography description and the presence, absence and count of the seven study species. Voucher specimens were collected for subsequent analysis, and are deposited in the Oman Botanic Garden herbarium (OBG) (Table 1). Summary statistics of quadrats were calculated in in R software V3.1.2, implemented in RStudio (R Development Core Team, 2014; RStudio Team, 2015). A checklist of fog moisture capture and water use efficiency functional traits commonly observed in xerophytic plants was also collated, following the approach of (Larraín-Barrios *et al.* 2018). Each study species was examined and scored for their presence/absence and degree of development (see Tables S1 and S2, Supporting Information). In addition to observations recorded during this field study, historical records were included from relevant national and international collections, specifically; Oman Botanic Garden Herbarium (OBG), Sultan Qaboos University (SQUH), the Oman Natural History Museum National Herbarium (ON) and the Royal Botanic Garden Edinburgh (E) (Summarised in Table 2; Figure S1, Supporting Information).

**Table 2.** Environmental niche model input and evaluation statistics.

| Species | Pre-filtering observations |  |  | Post-filtering | Model evaluation |  |  |  |  |  |
| --- | --- | --- | --- | --- | --- | --- | --- | --- | --- | --- |
|  | Incidental | Quadrats | Historical |  | Features | RM | AUC<sub>TEST</sub> | AUC<sub>DIFF</sub> | OR<sub>MTP</sub> | Parameters |
| AA | 161 | 20 | 12 | 41 | L | 2.5 | 0.84 | 0.11 | 0.05 | 6 |
| HG | 34 | 4 | 19 | 23 | L | 1.5 | 0.82 | 0.08 | 0.04 | 5 |
| OH | 16 | 3 | 13 | 13 | L | 0.5 | 0.79 | 0.15 | 0.23 | 6 |
| PJ | 8 | 0 | 1 | 4 | - | - | - | - | - | - |
| PP | 19 | 45 | 18 | 54 | LQHP | 2.5 | 0.76 | 0.11 | 0.04 | 10 |
| SG | 35 | 7 | 0 | 17 | L | 2 | 0.93 | 0.03 | 0.12 | 4 |
| SH | 15 | 15 | 0 | 25 | LQ | 1.5 | 0.89 | 0.05 | 0.04 | 8 |

Feature classes: Linear L, Quadratic Q, Hinge H, Product P and Threshold T. RM = regularization multiplier.

### Preparation of environmental variables

To ensure we captured important environmental variation, we collated 54 bioclimatic variables covering the study area at 1 km resolution (Table S3, Supporting Information). In addition to WorldClim2 and Bioclim variables (Fick and Hijmans 2017), we generated a complementary set of bioclimatic layers that may better characterise arid environments using the ‘ENVIREM’ package (Title and Bemmels 2018). We also sought to include night time fog, an important candidate variable in determining plant distributions in this region (Price et al. 1988). Specifically we followed the approach of Chaurasia *et al.* (2011) and classified fog based on the brightness temperature difference (ΔBT) of the 3.9 and 10.75 µm bands (channels 22 and 31) of the MODIS satellite. The emissive properties of these two bands differ for fog water droplets which are typically small, and do not excite the 3.9 µm band, whereas emissivity for both cloud and fog droplets is approximately the same for the 10.75 µm band (Hunt 1973). Twice nightly images at 1 km resolution were collated for the period 2001-17 (MODIS product: MOD021KM) from the LAADS database (MCST, 2018). Raw radiance values were converted to brightness temperature using Planck’s function implemented in ENVI software (Harris Geospatial) and the difference calculated. In contrast to Chaurasia *et al.* (2011), high quality real-time ground truth data is not available for our region, therefore we did not apply a fog classification threshold, instead we retained the data as a continuous variable with higher (less negative) values considered more likely to represent smaller fog water droplets.

At a fine spatial scale, other variables may also interact with fog moisture and influence the local ecology (Rastogi *et al.* 2016; Chung *et al.* 2017), thus we also incorporated several relevant fog proxies or co-variates. Cloud cover data (period 2001-15) was extracted from the global high-resolution cloud cover dataset generated by Wilson & Jetz (2016). Roughness, Terrain Ruggedness Index (TRI), slope and aspect were generated from a digital elevation model (GTOPO30) using the R package ‘Raster’ (Hijmans, 2017). Night time Land Surface Temperature (LST) was obtained from the MODIS satellite mission at 1 km resolution (Wan et al., 2018). Important climate variables are plotted, together with the study, area using the package ‘RasterVis’ (Lamigueiro, 2018).

To compare and characterize the range of environmental conditions across the Central Desert and other regional centres of endemism we randomly sampled all environmental variables for *n* cells in each region (with *n* being proportional to the area of the sampled polygon) and performed principal components analysis (PCA). We report variable loadings of the first and second principal components in Figure S2 (Supporting Information). To provide an additional line of evidence for local climatic stability, we compared interpolated Worldclim data (mean for 1970-2000) to more recent independent meteorological records from four contemporary weather stations (data period 1999-2017) in the central desert. Additional mapping and data visualization was performed using ‘ggplot2’ (Wickham 2009) and ‘rgeos’ (Bivand *et al.* 2018).

### Environmental niche modelling

The suite of environmental layers retained for modelling was refined in three stages. First, layers that had low variability at the spatial scale of our study area were removed (e.g. soil). Second, correlated environmental variables across the study area (r > |0.7|) were grouped, with a single variable from each group considered to be most relevant to arid environment plant ecology retained. Third, we performed an iterative selection procedure by removing variables with the highest Variance Inflation Factor (VIF) using the package ‘usdm’ (Naimi *et al.* 2014), with an upper threshold of VIF ≤ 2.5. Retained environmental variables are reported in Table S3 (Supporting Information).

Environmental niche modelling was performed with MAXENT v3.3.3 (Phillips et al., 2006), implemented in the packages ‘Dismo’ (Hijmans et al., 2011) and ‘ENMeval’ (Muscarella *et al.* 2014). To minimise model over-fitting, species data (including historic and incidental observations) were geographically rarefied to a 3 km bin size and examined across environmental space. Models were individually run and tuned for each study species over a study area encompassing the southern system, with quadrat surveys providing both presence and true absence data. Due to low sample sizes, data were partitioned using a jackknife approach where the number of model runs is equal to the number of occurrence localities, with a single data point excluded from each run for testing. Runs were performed iteratively across the full range of feature classes, with regularization multiplier values increasing from 1 to 4 in 0.5 increments.

Models were evaluated based on Akaike’s Information Criterion corrected for small sample sizes (i.e. (ΔAICc = 0), which penalises models that employ a greater number of parameters to describe the data (Warren and Seifert 2011; Muscarella *et al.* 2014). We report AUC_TEST_ averaged over all iterations, with higher values reflecting better model discrimination of presence locations from background absences. To quantify model overfitting we use two metrics. First, we report the mean difference in AUC between training and test data (AUC_DIFF_); this is expected to be higher where models are overfit to training data (Muscarella *et al.* 2014). Second, we report the proportion of testing localities with predicted habitat suitability values lower than the training locality with the lowest reported value (OR_MTP_). For each species, the best performing model was projected across the study area and a Maximum Training Sensitivity Plus Specificity (MaSS) logistic threshold, which balances the trade-off between omission and commission errors (Lobo *et al.* 2008; Liu *et al.* 2016), was employed to estimate habitat area. For two species (OH, SG) where model evaluation indicated evidence of weak overfitting, we used a Minimum Training Presence (MTP) threshold to ensure that all training observations are included within the predicted suitable habitat area.

In an effort to understand the abiotic drivers of the resulting distributions, several studies have shown that ranking variable contributions successfully captures biologically important factors (Kearney and Porter 2009; Searcy and Shaffer 2016). To assess relative variable importance across species we compare ranked permutation importance using Kendall’s *W*, corrected for ties, implemented in the package ‘irr’ (Gamer *et al.* 2012). Secondly, we use linear regression to assess the relationship between the contribution of mean annual fog to model performance, and the species’ trait score (see Tables S1-2, Supporting Information). Finally we calculated niche overlap across study species using the method of Warren, Glor and Turelli, (2008) and then combined thresholded species distribution classifications to identify areas of spatial overlap and co-occurrence of multiple species. RBG images were obtained from Sentinel 2 (Copernicus Sentinel data 2015, processed by ESA, accessed from https://remotepixel.ca/ on 20/12/2018) and plotted with increased contrast. Surface wind direction data, averaged for the months June to August (2015-17), was obtained from the Global Forecasting System, via the package ‘rWind’ (Fernández-López 2018).

## RESULTS

Evaluation of regional climate identifies a weak influence of the southern monsoon system on the Jiddat Al Arkad of the southern system (Figure 2). Concurrently, increased summer fog incidence in the southern Central Desert coincides with the warmest temperatures of the summer months, which appears to result in cooler coastal night time temperatures. Principal component analysis of abiotic variables clearly differentiated the major regions of endemism (Figure 3). Overall temperature related variables were the major contributors to PC1, with precipitation differentiating PC2 (Figure S2, Supporting Information). The two Central Desert systems are found to be differentiated, but partially overlapping with 88.5% and 62% of points representing a unique climate variables for the northern and southern systems respectively.

**Figure 2.**
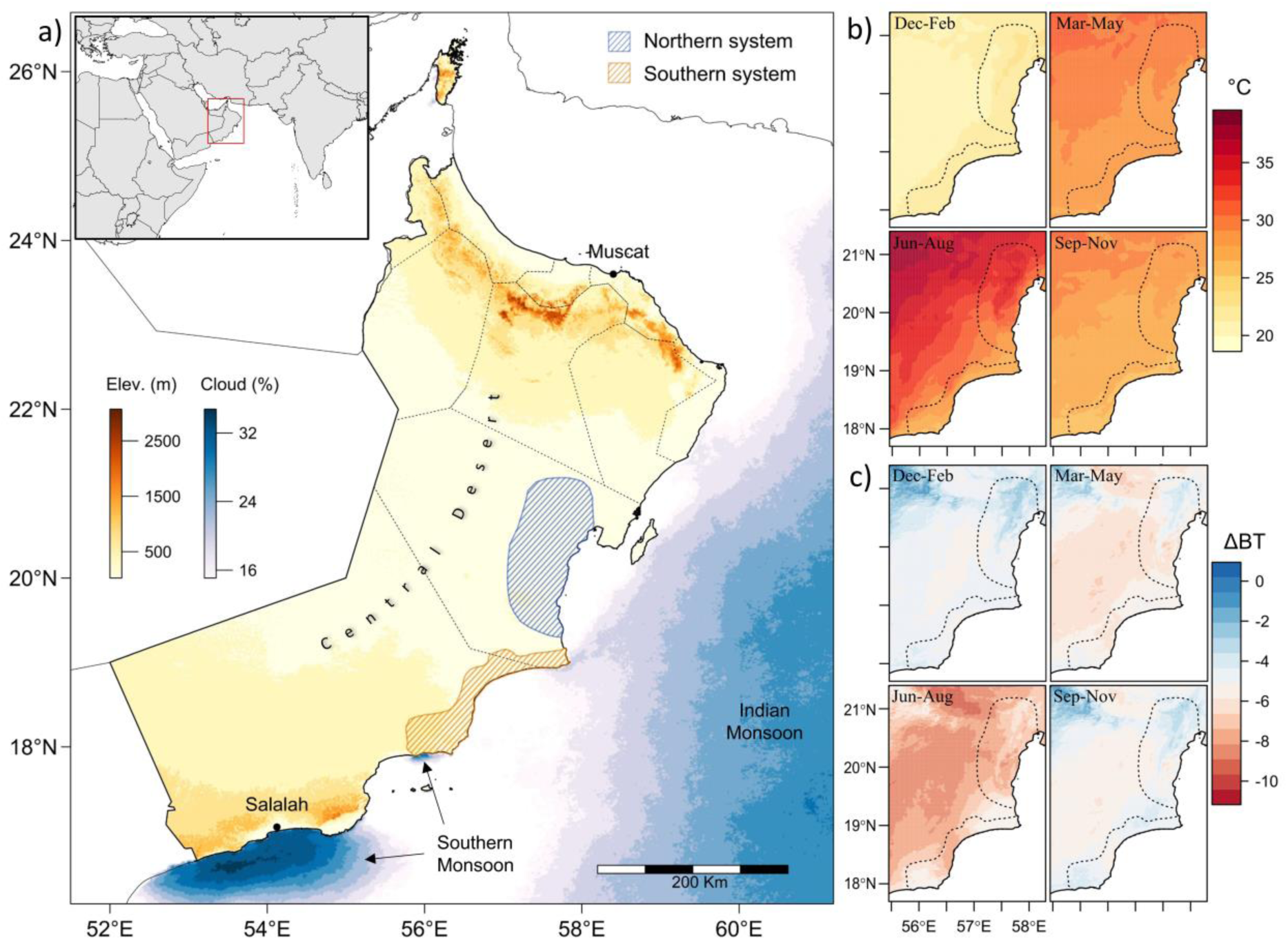
a) Elevation map of Oman, with annual offshore cloud cover percentage identifying the major southern and Indian monsoon climate patterns. Cloud cover over land is not shown, but is negligible for the Central Desert. Northern and southern study systems are denoted by shaded polygons. b) Quarterly mean temperature across the Central Desert. c) Quarterly night time fog intensity (change in brightness temperature) across the Central Desert. Higher values (less negative) are indicative of greater fog intensity.

**Figure 3.**
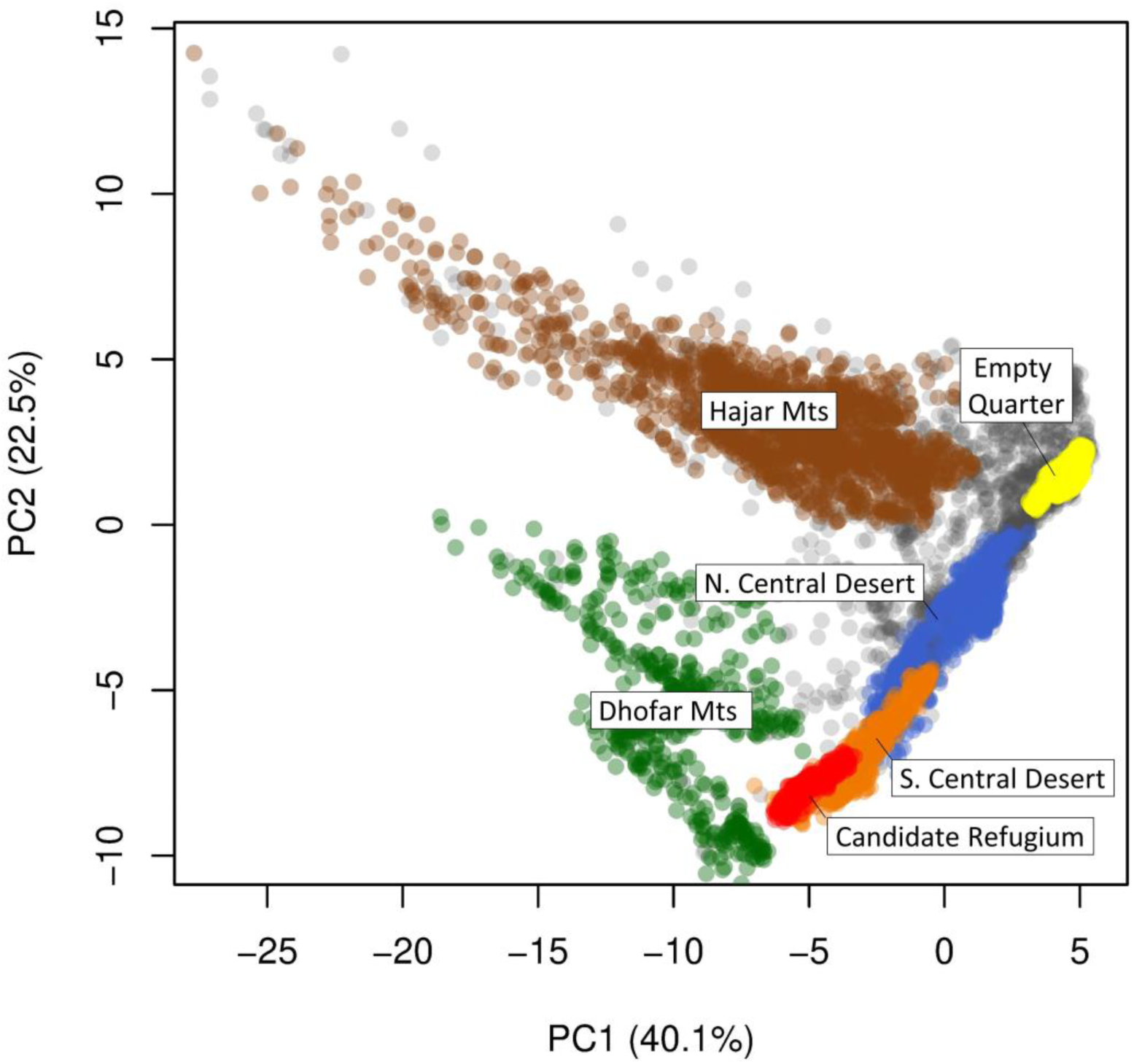
Principal component analysis of abiotic variables across principal ecoregions of Oman. Grey points denote a random background sample from across Oman. The five major centres of plant endemism comprise i) the Hajar Mountains; ii-iii) the Dhofar Mountains comprising the Jabal Samhan and Jabal Al Qamar/Qara centres of endemism, combined for the purposes of this figure; iv) the Northern Central Desert consisting of the Jiddat Al Harasis/Huqf and v) the Southern Central Desert comprising the Sahil Al Jazir/Jiddat Al Arkad. The Empty Quarter is plotted (yellow) for reference as it comprises a significant proportion of Oman’s land area, but is not considered a centre of endemism. The candidate refugium (red) is a subset of the southern system of the Central Desert.

Fieldwork surveys assessed 195 quadrats throughout the Southern System and successfully located all seven regional endemics. Study species were recorded in 41% of quadrats. In addition, 288 incidental and 68 historical observations were collated. After spatial filtering, 177 records were retained (Table 2; Figure S1, Supporting Information). Mean pH and EC across all quadrats was 7.73 (SD = 0.44) and 14.99 (SD = 27.13) respectively. No significant difference in pH or EC values was identified across species (ANOVA p > 0.05). In our assessment of fog and drought functional trait adaptation, *P. pulvinata* scored highest, and *S. gallagheri* scored lowest, with low stature, low leaf area and hairs the most frequent adaptations.

### Environmental niche modelling and variable importance

After filtering, we retained nine environmental variables for modelling (Table S3, Supporting Information). Ecological niche model evaluation statistics are reported in Table 2. Modelling was not performed for *P. jazirensis* due to insufficient data. AUC_TEST_ ranged from 0.76 (PP) to 0.93 (SG). AUC values are often lower for more widespread species, which may be the case for *P. pulvinata* and *O. Harsusiticus* (Jiménez-Valverde *et al.* 2008). Model logistic habitat suitability projections are plotted in Figure 4, with the percentage contribution of variables reported in Table 3. Binary threshold maps are provided in Figure S3 (Supporting Information). The most important variables varied substantially, with no evidence of consistent rank importance across species (*Wt* = 0.079, p = 0.87). Annual mean fog did not appear to rank highly for any species, and was not significantly associated with fog adaptation trait scores (F_1,4_ = 0.76, p = 0.4). Niche overlap was high in all pairwise comparisons (median = 0.87; Table S4, Supplementary Information).

**Table 3.** Percentage contribution of environmental variables to environmental niche models across study species.

| Environmental variable | AA | HG | OH | PP | RG | SH | Mean (SD) |
| --- | --- | --- | --- | --- | --- | --- | --- |
| Night LST | 22.7 | 0.0 | 0.0 | 1.3 | 4.1 | 2.6 | 5.1 (8.8) |
| Mean annual cloud cover | 0.0 | 2.2 | 0.0 | 2.4 | 0.0 | 1.6 | 1.0 (1.2) |
| Cloud cover seasonality | 0.5 | 31.2 | 0.0 | 2.9 | 0.4 | 9.4 | 7.4 (12.2) |
| Thornthwaite aridity index | 44.5 | 0.0 | 4.2 | 0.0 | 82.3 | 0.8 | 22.0 (34.3) |
| Aspect | 0.0 | 0.0 | 10.2 | 1.6 | 0.0 | 33.7 | 7.6 (13.4) |
| Elevation | 0.0 | 40.6 | 33.1 | 37.7 | 0.0 | 26.7 | 23.0 (18.4) |
| Mean annual fog | 8.0 | 3.4 | 13.7 | 2.5 | 0.9 | 9.0 | 6.2 (4.8) |
| Terrain roughness index | 24.0 | 0.0 | 7.6 | 21.1 | 12.4 | 5.0 | 11.7 (9.4) |
| Annual mean temperature | 0.5 | 22.6 | 31.3 | 30.4 | 0.0 | 11.1 | 16.0 (14.2) |

**Figure 4.**
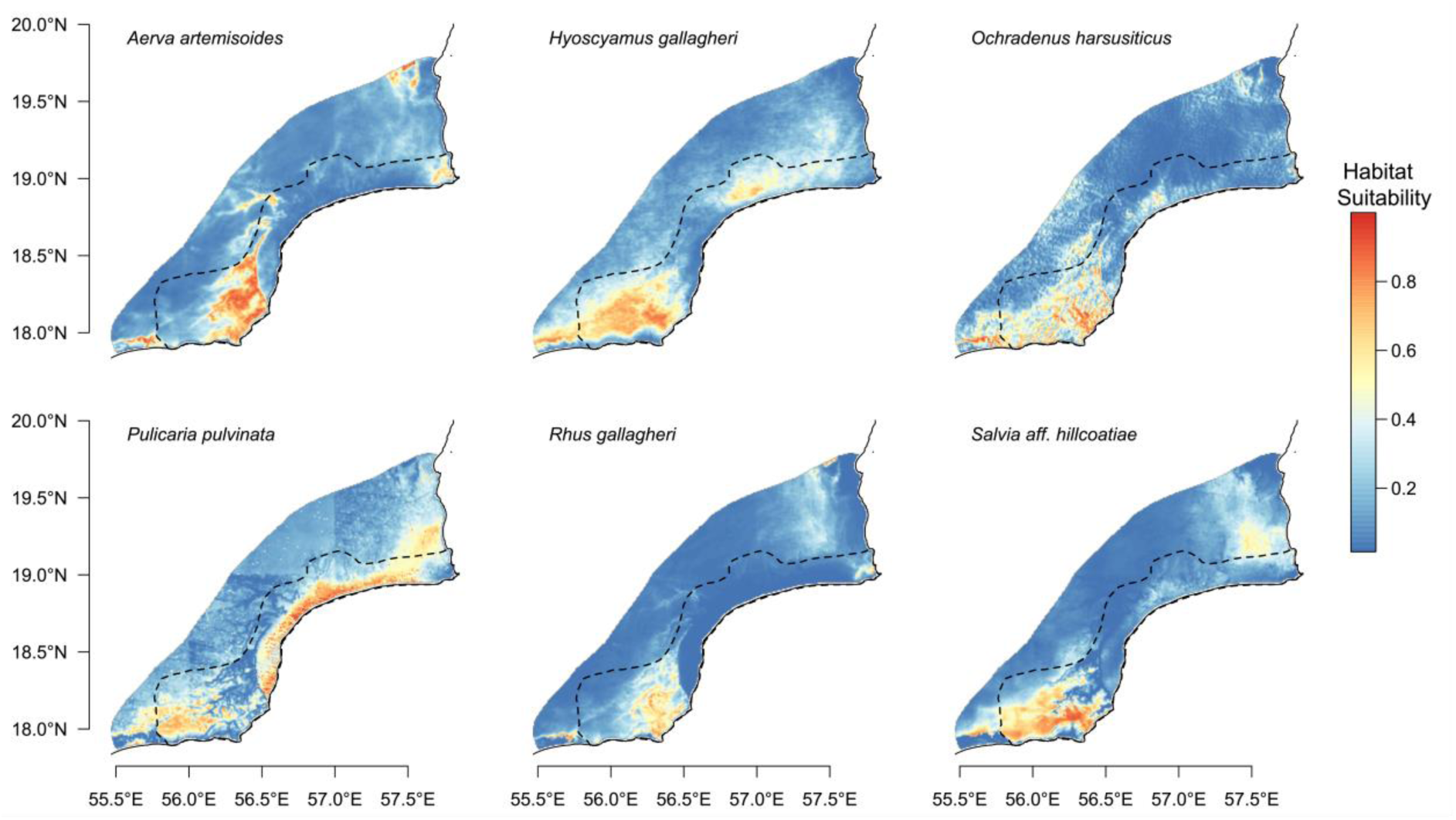
Environmental niche models for each of six study species across the southern Central Desert. Higher values indicative of greater modelled habitat suitability. Dashed line denotes the southern Central Desert system.

### Identification of climate refugia

Combined binary species distributions identified a key area where all study species are predicted to co-occur (Figure 5A). High predicted habitat suitability across all models was localized to the southern Jiddat al Arkad. Satellite imagery shows the region, seasonal cloud cover and the prevailing summer wind direction in Figure 5B. Independent contemporary weather station records provide an additional line of evidence. Whilst the three northern stations show elevated maximum daily temperatures (period 2002-17) compared to the Worldclim 2 reference (1970-2000), Shalim station – close to our putative coastal fog and cloud affected area – shows summer maximum temperatures below the Worldclim 2 reference (Figure 6).

**Figure 5.**
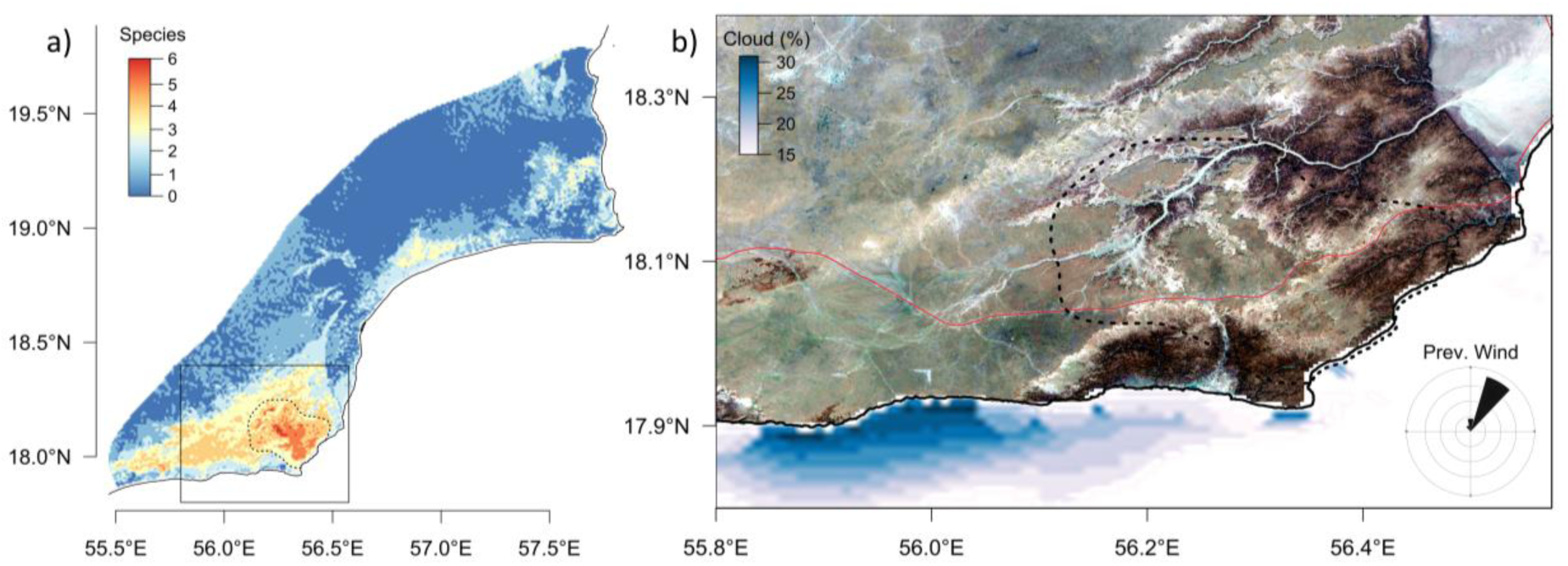
Identification of endemic species co-occurrence in the southern Central Desert. a) Composite map of the binary distributions of six study species. Dashed line identifies a region of high diversity with potential as a candidate Important Plant Area. b) False colour Sentinel 2 image of the high diversity area. Red line shows a primary road crossing the study area. Inset rose diagram shows the prevailing wind direction during the Khareef (Southern Monsoon). Cloud cover mean is shown in blue.

**Figure 6.**
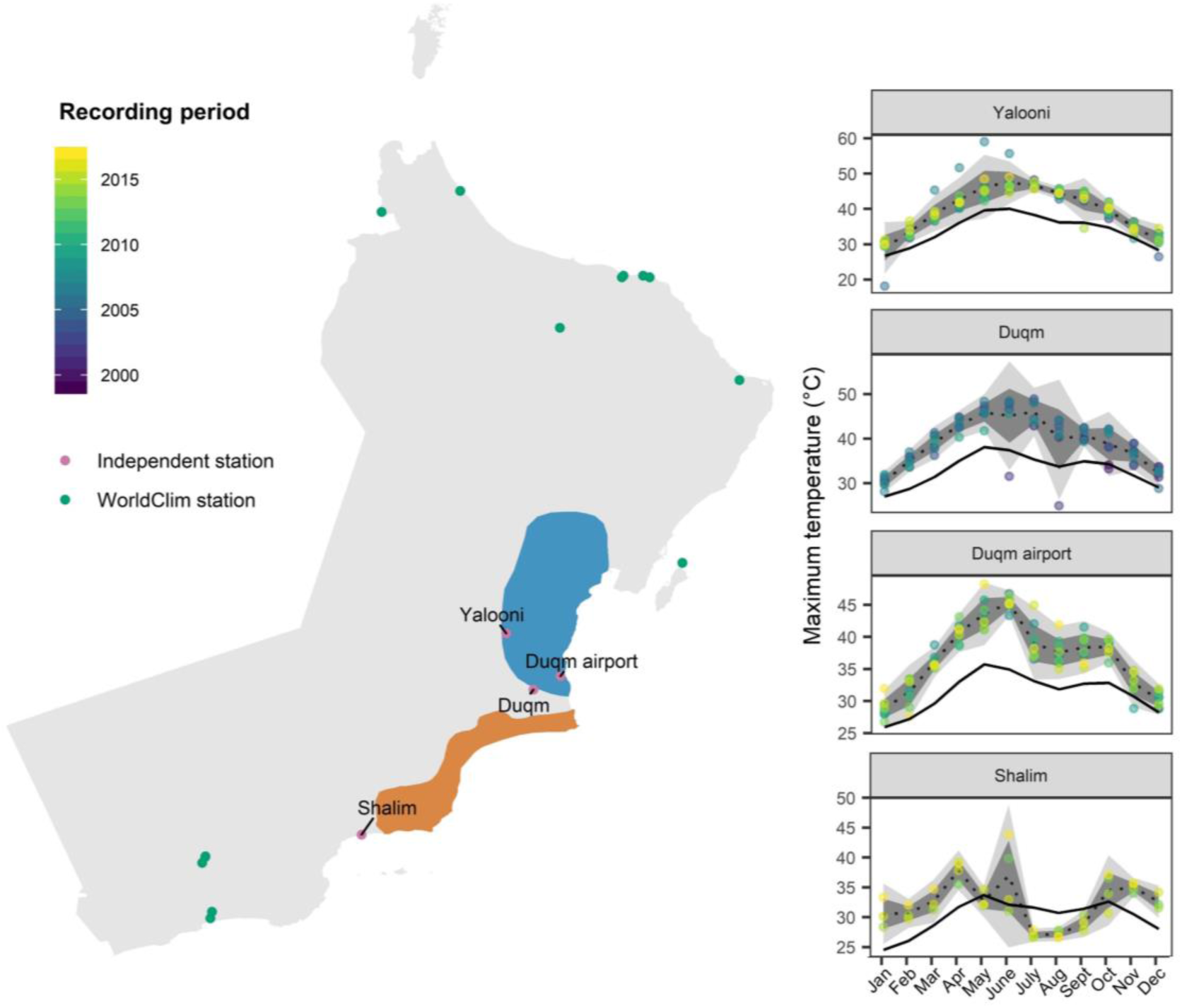
Locations of Omani weather stations contributing to interpolated climate variables used in this study (1970-2000), as well as four independent validation weather stations from the Central Desert (1999-2017). Maximum daily temperature recorded at these stations is reported (coloured by year), with the black line denoting the historic Worldclim 2 average for this period. Northern and southern systems are depicted in blue and orange, respectively.

## DISCUSSION

In this study we present evidence for a Southern Arabian Pleistocene refugium in Oman’s Central Desert (Figures 2, 5). As shown by Sandel et al. (2011), the negative relationship between endemism and the increasing velocity of changing climate is strongest in poorly dispersing species such as plants. Therefore co-occurrence of a high number of endemic study species within a narrow monsoon-influenced region is indicative of a refugium with low climate change velocity (Sandel *et al.* 2011; Abellán and Svenning 2014; Harrison and Noss 2017). Climate analysis identified cooler mean annual temperatures in the study area and the influence of coastal cloud and fog (Figure 2), which combined with novel survey data and environmental distribution modelling suggests that the vegetation of the southern Central Desert is a relict of an earlier, more mesic period. This is further supported by the biogeography of genera such as *Aerva*, *Searsia* and *Ochradenus* which have global distributions from Africa to South-East Asia, yet with endemic species restricted to Arabia (POWO 2018), indicating support for a refugium further back into the Neogene. Thus, this system may represent the northernmost remnant of a continuous belt of mesic vegetation formerly ranging from Africa to Asia, with close links to the flora of East Africa (Kürschner, 1998; Patzelt, 2011).

The relictual distributions that we observe appear to be driven by the interaction of climate and topographic factors, in particular the influence of the southern monsoon. It had been speculated that parts of the Central Desert may be at the fringe of the monsoon-affected area (Patzelt, 2015), thus benefiting from occasional low clouds, cool winds and coastal fog during the southern monsoon, but previously this could not be tested because of the lack of climate stations. Here, using evidence from remote sensing, we demonstrate that the southern monsoon does indeed influence the southern system of the Central Desert, with patterns of night time fog detected via the MODIS satellite also consistent with the limited reports available. This putatively places the southern Arabian coastal fog-influenced Central Desert together with other coastal fog deserts such as the Namib and Peruvian lomas (Cereceda *et al.* 2008; Henschel and Seely 2008), though based on limited fog adaptation traits in the flora, fog intensity may be lower.

By applying these climate data to systematic field records of endemic plants, we develop a suite of models characterizing each species’ environmental niche. We show that despite small sample sizes it is possible to generate robust niche models, incorporating true absence data, which identify important areas of plant diversity. Surprisingly, the relative importance of retained climatic and topographic variables differed substantially across study species. Therefore, we conclude that it is not a single set of environmental variables contributing to the distribution of this unique flora. For example, whilst overall, aridity and mean annual temperature are unsurprisingly important predictors in an arid environment, almost all retained variables are important across specific taxa. Therefore our analysis does not support the hypothesis that it is predominantly fog that influences the distribution of this endemic flora, rather a range of factors appear to be important, consistent with the diverse traits and phylogenetic provenance of the species. We note however that on a finer spatial scale, factors such as microrefugia and fog hydrology may have greater importance (Mclaughlin *et al.* 2017).

Despite being recognised for its global importance, the arid Horn of Africa biodiversity hotspot is one of the most severely degraded, with less than 5% of habitat considered to be in pristine condition (Mittermeier *et al.* 2005; Mallon 2013). Key threats to the Central Desert include overgrazing (Ghazanfar, 2004), and climate change (Almazroui *et al.* 2013), with mean annual temperature for the Arabian Peninsula increasing at 0.6°C per decade and a significant decreasing trend in annual rainfall (Almazroui *et al.* 2013). It is also concerning that climate change has been associated with a historic shifts in intensity and northward extent of the monsoon (Fleitmann and Matter 2009) and elsewhere a contemporary decline in coastal fog frequency (Johnstone and Dawson 2010), with strong implications for persistence of endemic flora. In our study area, a relatively minor shift in the northward extent of the monsoon could have significant implications for regional climate.

Refugia have been suggested as priority sites to conserve global biodiversity under climate change precisely because of their demonstrated ability to facilitate species survival under adverse conditions (Keppel *et al.* 2012). Based on previous studies, it is also likely that refugial populations harbour the highest genetic diversity across the species’ distribution (Meister, Hubaishan, Kilian, & Oberprieler, 2005), helping building future evolutionary resilience (Sgrò *et al.* 2011). This may be particularly important in the Central Desert, where many endemic species have been restricted to only a single refugial location, reducing potential for subsequent population admixture (Petit *et al.* 2003). The southern coast of the Arabian peninsula has also been predicted to contain a significant proportion of unassessed at-risk vascular plant species (Pelletier *et al.* 2018). In recent years, an Important Plant Area (IPA) programme has been initiated for the Middle East, which highlights the value of an ecological and evolutionary process-based view in identifying candidate conservation sites (Al-Abbasi *et al.* 2010).

Therefore, the site outlined here (Figure 5), covering approximately 880 km^2^, may be candidate for further evaluation and consideration as an IPA.

In conclusion, this study makes an important contribution to our understanding of southern Arabian climate refugia, and the biogeographical origins of the endemic flora of Oman’s Central Desert. In the future we highlight the value of a network of detectors to characterise coastal fog across the landscape, particularly in the southern Central Desert. These would better enable an assessment of how coastal fog co-varies with other readily available datasets such as topography, cloud cover and land surface temperature to enable higher resolution predictions of the influence of coastal fog on species distributions. More generally, we emphasise the value of predictive modelling in the region to advance beyond initial presence-absence grids, both to identify the drivers of biogeographic patterns and to prioritise sites for the conservation. In the future, further identification and characterisation of southern Arabian climate refugia may be a useful strategy to support conservation in a global biodiversity hotspot.

## Supporting information

Borrell Supplementary Materials

## ACKNOWLEDGEMENTS

We gratefully acknowledge field support from Fathi Al Hasani and Salim Al Rahbi, and a team of UK volunteers who assisted in data collection and logistics.

## FUNDING INFORMATION

This work was supported by a fieldwork grant from the Anglo-Omani Society, UK, to the Central Desert Botanic Expedition team.

## DATA ACCESSIBILITY

All topographic and environmental GIS layers used in this study are freely available from the sources outlined in Table S3, Supporting Information. Raw meteorological data for Central Desert climate stations are available on request from the Meteorological Society of Oman. Novel species observation records from the Central Desert will be provided on the Global Biodiversity Information Facility.

## BIOSKETCH

The research reported here emerged from the Central Desert Botanic Expedition 2017 and is the result of an ongoing collaboration between British scientists, volunteer participants and the Oman Botanic Garden. Initiated in collaboration with the British Exploring Society in 2012, collaborative teams have conducted research across several locations in Dhofar with further research planned in the Musandam peninsula. Fieldwork has been generously supported by the Anglo-Omani Society which seeks to promote understanding and friendship between Britain and Oman, particularly through scientific and cultural exchange. J.B., G.I., D.L., A.S.R. and A.P. conceived the idea, all authors (bar T.W. and A.P.) participated in fieldwork. J.B., T.S., R.S. and T.W. performed analysis and J.B., D.L. and A.P. led the writing. All authors approved the final version of this manuscript.

## LITERATURE CITED

Abellán P, Svenning JC. 2014. Refugia within refugia - patterns in endemism and genetic divergence are linked to Late Quaternary climate stability in the Iberian Peninsula. Biological Journal of the Linnean Society 113: 13–28.

Al-Abbasi TM, Al-Farhan A, Al-Khulaidi AW, et al. 2010. Important plant areas in the arabian peninsula. Edinburgh Journal of Botany 67: 25–35.

Almazroui M, Abid MA, Athar H, Islam MN, Ehsan MA. 2013. Interannual variability of rainfall over the Arabian Peninsula using the IPCC AR4 global climate models. International Journal of Climatology 33: 2328–2340.

Bennett KD, Tzedakis PD, Willis KJ. 1991. Quaternary refugia of north European trees.: 103–115.

Birks HJB, Willis KJ. 2008. Alpines, trees, and refugia in Europe. Plant Ecology and Diversity 1: 147–160.

Bivand R, Stuetz R, Ove K, Giraudoux P, Davis M, Santilli S. 2018. Package ‘ rgeos.’

Brinkmann K, Dickhoefer U, Schlecht E, Buerkert A. 2011. Quantification of aboveground rangeland productivity and anthropogenic degradation on the Arabian Peninsula using Landsat imagery and field inventory data. Remote Sensing of Environment 115: 465–474.

Cereceda P, Larrain H, Osses P, Farías M, Egaña I. 2008. The spatial and temporal variability of fog and its relation to fog oases in the Atacama Desert, Chile. Atmospheric Research 87: 312–323.

Chaurasia S, Sathiyamoorthy V, Paul Shukla B, Simon B, Joshi PC, Pal PK. 2011. Night time fog detection using MODIS data over Northern India. Meteorological Applications 18:483–494.

Chung M, Dufour A, Pluche R, Thompson S. 2017. How much does dry-season fog matter? Quantifying fog contributions to water balance in a coastal California watershed. Hydrological Processes 31: 3948–3961.

Comes HP, Kadereit JW. 1998. The effect of quaternary climatic changes on plant distribution and evolution. Trends in Plant Science 3: 432–438.

Delany MJ. 1989. The zoogeography of the mammal fauna of southern Arabia. Mammal Review 19: 133–152.

Dilts TE, Weisberg PJ, Dencker CM, Chambers JC. 2015. Functionally relevant climate variables for arid lands: A climatic water deficit approach for modelling desert shrub distributions. Journal of Biogeography 42: 1986–1997.

Dregne HE. 1976. Soils of Arid Regions. Amsterdam: Elsevier Scientific Publishing Company.

Elith J, Leathwick J. 2009. Species distribution models: ecological explanation and prediction across space and time. Annual Review of Ecology, Evolution, … 40: 677–697.

Fernández-López J. 2018. rWind: Download, edit and include wind data in ecological and evolutionary analysis. Ecography 42.

Fick SE, Hijmans RJ. 2017. Fick, Hijmans - 2017 - WorldClim 2 new 1-km spatial resolution climate surfaces for global land areas.pdf.

Fischer DT, Still CJ, Williams AP. 2009. Significance of summer fog and overcast for drought stress and ecological functioning of coastal California endemic plant species. Journal of Biogeography 36: 783–799.

Fisher M, Membery DA. 1998. Climate In: Ghazanfar SA, Fisher M, eds. Vegetation of the Arabian Peninsula. Dordrecht, The Netherlands: Kluwer Academic Publishers, 5–38.

Fleitmann D, Matter A. 2009. The speleothem record of climate variability in Southern Arabia. Comptes Rendus - Geoscience 341: 633–642.

Gamer M, Lemon J, Fellows I, Singh P. 2012. Irr package for R, version 0.84.

Gandini F, Achilli A, Pala M, et al. 2010. Mapping human dispersals into the Horn of Africa from Arabian Ice Age refugia using mitogenomes. Scientific Reports 6: 1–13.

Ghazanfar SA. 1998. Status of the flora and plant conservation in the sultanate of Oman. Biological Conservation 85: 287–295.

Ghazanfar SA. 2004. Biology of the central desert of Oman. Turkish Journal of Botany 28: 65–71.

Ghazanfar SA, Fisher M. 2013. Vegetation of the Arabian peninsula. Springer Science & Business Media.

Ghazanfar S, Gallagher M. 1998. Remarkable lichens from the Sultanate of Oman.

Harrison S, Noss R. 2017. Endemism hotspots are linked to stable climatic refugia. Annals of Botany 119: 207–214.

Henschel JR, Seely MK. 2008. Ecophysiology of atmospheric moisture in the Namib Desert. Atmospheric Research 87: 362–368.

Hijmans RJ. 2017. raster: Geographic Data Analysis and Modeling. R package version 2.6–7.

Hijmans RJ, Philips S, Leathwick J, Elith J. 2017. dismo: Species Distribution Modeling. R package version 1.1–4.

Hunt GE. 1973. Radiative properties of terrestrial clouds at visible and infra-red thermal window wavelengths. Quarterly Journal of the Royal Meteorological Society 99: 346–369.

Jennings RP, Singarayer J, Stone EJ, et al. 2015. The greening of Arabia: Multiple opportunities for human occupation of the Arabian Peninsula during the Late Pleistocene inferred from an ensemble of climate model simulations. Quaternary International 382: 181–199.

Jiménez-Valverde A, Lobo JM, Hortal J. 2008. Not as good as they seem: The importance of concepts in species distribution modelling. Diversity and Distributions 14: 885–890.

Johnstone JA, Dawson TE. 2010. Climatic context and ecological implications of summer fog decline in the coast redwood region James. Proceedings of the National Academy of Sciences 107: 4533–4538.

Jolly D, Prentice IC, Bonnefille R, et al. 2009. Biome Reconstruction from Pollen and Plant Macrofossil Data for Africa and the Arabian Peninsula at 0 and 6000 Years Published by: Blackwell Publishing Stable URL: http://www.jstor.org/stable/2846197.

Kearney M, Porter W. 2009. Mechanistic niche modelling: Combining physiological and spatial data to predict species’ ranges. Ecology Letters 12: 334–350.

Keppel G, Van Niel KP, Wardell-Johnson GW, et al. 2012. Refugia: Identifying and understanding safe havens for biodiversity under climate change. Global Ecology and Biogeography 21: 393–404.

Kürschner H. 1998. Biogeography and Introduction to Vegetation. In: Ghazanfar, S.A., Fisher M, ed. Vegetation of the Arabian Peninsula. Dordrecht: Springer,.

Larraín-Barrios B, Faúndez-Yancas L, Búrquez A. 2018. Plant functional trait structure in two fog deserts of America. Flora: Morphology, Distribution, Functional Ecology of Plants 243: 1–10.

Liu C, Newell G, White M. 2016. On the selection of thresholds for predicting species occurrence with presence-only data. Ecology and Evolution 6: 337–348.

Lobo JM, Jiménez-Valverde A, Real R. 2008. AUC: a misleading measure of the performance of predictive distribution models. Global Ecology and Biogeography 17: 145–151.

Mallon DP. 2013. Global hotspots in the Arabian Peninsula. Zoology in the Middle East 7140.

Mclaughlin BC, Ackerly DD, Klos PZ, Natali J, Dawson TE, Thompson S. 2017. Hydrologic refugia, plants, and climate change. Global Change Biology 23: 2941–2961.

Meister J, Hubaishan MA, Kilian N, Oberprieler C. 2005. Chloroplast DNA variation in the shrub Justicia areysiana (Acanthaceae) endemic to the monsoon affected coastal mountains of the southern Arabian Peninsula. Botanical Journal of the Linnean Society 148: 437–444.

Meister J, Hubaishan MA, Kilian N, Oberprieler C. 2006. Temporal and spatial diversification of the shrub Justicia areysiana Deflers (Acanthaceae) endemic to the monsoon affected coastal mountains of the southern Arabian Peninsula. Plant Systematics and Evolution 262: 153–171.

Merow C, Smith MJ, Silander JA. 2013. A practical guide to MaxEnt for modeling species’ distributions: What it does, and why inputs and settings matter. Ecography 36: 1058–1069.

Miller AG. 1994. Dhofar Fog Oasis, Oman and Yemen. In: Davis SD, Heywood VH, Hamilton A., eds. Centres of Plant Diversity, Vol. 1. Gland: WWF, IUCN., 143–155.

Miller AG, Cope TA. 1996. Flora of the Arabian Peninsula and Socotra, 1. Edinburgh: Edinburgh University Press.

Miller AG, Nyberg JA. 1990. Patterns of endemism in Arabia In: Contributiones selectae ad floram et vegetationem orientis: proceedings of the Third Plant Life of southwest Asia Symposium, held.3–8.

Mittermeier RA, Gil PR, Hoffman M, et al. 2005. Hotspots Revisited: Earth’s Biologically Richest and Most Endangered Terrestrial Ecoregions. Monterrey, Mexico: Cemex, Conservation International and Agrupación Sierra Madre.

MODIS Characterization Support Team (MCST). 2018. MODIS 1km Calibrated Radiances Product.

Muscarella R, Galante PJ, Soley-Guardia M, et al. 2014. ENMeval: An R package for conducting spatially independent evaluations and estimating optimal model complexity for Maxent ecological niche models. Methods in Ecology and Evolution 5: 1198–1205.

Naimi B, Hamm NAS, Groen TA, Skidmore AK, Toxopeus AG. 2014. Where is positional uncertainty a problem for species distribution modelling? Ecography 37: 191–203.

Parker AG. 2010. Pleistocene Climate Change in Arabia: Developing a Framework for Hominin Dispersal over the Last 350 ka.

Patel. 1992. Geological Map of Juzor Al Halaaniyaat, Sheet NE 40 - 10 and Duqm and Madraca Sheet NE 40 - 03/07.

Patzelt A. 2009. The mountain vegetation of northern Oman: Ecology, phytosociology and biogeography of Olea europaea and Juniperus excelsa woodlands and of weed vegetation on cultivated terraces. In: Victor, R., Robinson, M. (eds.) Al Jabal Al Akhdar Monograph.

Patzelt A. 2011. The themeda quadrivalvis tall-grass savannah of Oman at the crossroad between Africa and Asia. Edinburgh Journal of Botany 68: 301–319.

Patzelt A. 2014. Oman Plant Red Data Book. Diwan of Royal Court, Oman Botanic Garden.

Patzelt A. 2015. Synopsis of the Flora and Vegetation of Oman, with Special Emphasis on Patterns of Plant Endemism. Abhandlungen der Braunschweigischen Wissenschaftlichen Gesellschaft: 282–317.

Pelletier TA, Carstens BC, Tank DC, Sullivan J, Espíndola A. 2018. Predicting plant conservation priorities on a global scale. Proceedings of the National Academy of Sciences 115: 201804098.

Perpinan Lamigueiro O. 2018. Package “rasterVis” - Visualization Methods for Raster Data.

Peterson AT, Papeş M, Soberón J. 2015. Mechanistic and Correlative Models of Ecological Niches. European Journal of Ecology 1: 28–38.

Petit RJ, Petit J, Aguinagalde I, et al. 2003. Genetic Diversity Glacial Refugia: Hotspots But Not. Science 1563: 1563–5.

Phillips SJ, Anderson RP, Schapire RE. 2006. Maximum entropy modeling of species geographic distributions. Ecological Modelling 190: 231–259.

POWO. 2018. Plants of the World Online. http://www.plantsoftheworldonline.org/. 29 Dec. 2018.

R Development Core Team. 2014. R: A language and environment for statistical computing. Foundation for Statistical Computing, Vienna, Austria.

Rastogi B, Williams AP, Fischer DT, et al. 2016. Spatial and temporal patterns of cloud cover and fog inundation in coastal California: Ecological implications. Earth Interactions 20.

Raven PH, Axelrod DI. 1974. Angiosperm Biogeography and Past Continental Movements. Annals of the Missouri Botanical Garden 61: 539–673.

RStudio Team. 2016. RStudio: Integrated Development for R. RStudio, Inc., Boston, MA.

Sandel B, Arge L, Dalsgaard B, et al. 2011. The Influence of Late Quaternary. Science 334: 660–664.

Schemenauer RS, Cereceda P, Carvajal N. 1987. Measurements of Fog Water Deposition and Their Relationships to Terrain Features. Journal of Applied Meteorology 26: 1285–1292.

Searcy CA, Shaffer HB. 2016. Do Ecological Niche Models Accurately Identify Climatic Determinants of Species Ranges? The American Naturalist 187:4: 423–435.

Sgrò CM, Lowe AJ, Hoffmann AA. 2011. Building evolutionary resilience for conserving biodiversity under climate change. Evolutionary Applications 4: 326–337.

Stanley Price MR, Al-Harthy AH, Whitcombe RP. 1988. Fog moisture and its ecological effects in Oman In: Arid Lands: Today and Tomorrow.69–88.

Title PO, Bemmels JB. 2018. ENVIREM: an expanded set of bioclimatic and topographic variables increases flexibility and improves performance of ecological niche modeling. Ecography 41: 291–307.

Wan Z, Hook S, Hulley G. 2018. MOD11A1 MODIS/Terra Land Surface Temperature and the Emissivity Daily L3 Global 1km SIN Grid.

Wang N, Borrell JS, Bodles WJ a, Kuttapitiya A, Nichols R a., Buggs RJ a. 2014. Molecular footprints of the Holocene retreat of dwarf birch in Britain. Molecular Ecology 23: 2771–2782.

Warren DL, Glor RE, Turelli M. 2008. Environmental niche equivalency versus conservatism: Quantitative approaches to niche evolution. Evolution 62: 2868–2883.

Warren DL, Seifert SN. 2011. Ecological niche modeling in Maxent: the importance of model complexity and the performance of model selection criteria. Ecological Applications 21: 335–342.

White F, Léonard J. 1990. Phytogeographical links between Africa and southwest Asia In: Contributiones selectae ad floram et vegetationem orientis: proceedings of the Third Plant Life of southwest Asia Symposium, held.3–8.

Whybrow PJ, Mcclure HA. 1981. Fossil Mangrove Roots and Palaeoenvironments of the. 32: 213–225.

Wickham H. 2009. ggplot2. Elegant graphics for data analysis. Springer: 210.

Wilson AM, Jetz W. 2016. Remotely Sensed High-Resolution Global Cloud Dynamics for Predicting Ecosystem and Biodiversity Distributions. PLoS Biology 14: 1–20.

Zhang H, Schroder JL, Pittman JJ, Wang JJ, Payton ME. 2005. Soil Salinity Using Saturated Paste and 1:1 Soil to Water Extracts. Soil Science Society of America Journal 69: 1146.

