## Supplementary material for "Islands in the desert: Environmental distribution modelling of endemic flora reveals the extent of Pleistocene tropical relict vegetation in southern Arabia": Borrell Supplementary Materials

[1]Jodrell Laboratory, Royal Botanic Gardens, Kew, Richmond, Surrey, TW9 3DS, UK. [2]Oman Botanic Garden, Muscat, Oman. [3]RSPB Centre for Conservation Science, Royal Society for the Protection of Birds, The Lodge, Sandy, Bedfordshire, SG19 2DL, UK. [4]Department of Animal and Plant Sciences, Alfred Denny Building, University of Sheffield, Western Bank, Sheffield, S10 2TN. [5]Biodiversity Information Service, Unit 4, Royal Buildings, 6 Bulwark, Brecon, Powys, LD3 7LB. [6]Outward Bound Canada, 550 Bayview ave., Building 1 suite 201, Toronto, ON, M4W 3X8

**Running title**: A pleistocene refugium in southern Arabia

#### Supplementary Tables

**Table S1.** Functional traits associated with desert fog moisture capture and drought tolerance, adapted from Larraín-Barrios et al. (2018).

| Trait | Category | Weight | Adaptive relevance |
|---|---|---|---|
| **Plant morphology** | | | |
| Maximum plant height | 0 - 0.1 m | 1 | Greater height makes water stress cavitation more likely. |
| | 0.1 - 0.3 m | 0.5 | |
| | > 0.3 m | 0 | |
| Leaf surface area | <25 $mm^2$ | 1 | Smaller leaves are associated with reduced transpiration |
| | 25–225 $mm^2$ | 0.5 | |
| | >225 $mm^2$ | 0 | |
| Leaf and stem hairs | Pilose or tormentose | 1 | Higher density of hairs is associated with increased solar reflectance and reduced transpiration. |
| | Few, appressed or glandular hairs | 0.5 | |
| | Absent | 0 | |
| Succulence | Succulent leaves and stems | 1 | Water storage tissues to mitigate drought stress |
| | Succulent leaves | 0.5 | |
| | Succulent stems | 0.5 | |
| | Non-succulent | 0 | |
| Rosettes | Rosette | 1 | Leaf structure associated with increased fog interception |
| | Absent | 0 | |
| Photosynthetic stems | Photosynthetic (green) stem tissue | 1 | Associated with water use efficiency during drought |
| | Absent | 0 | |
| **Phenology** | | | |
| Phenology | Deciduous | 1 | Leaf loss associated with water use efficiency during drought stress |
| | Evergreen | 0 | |
| **Storage organs** | | | |
| Degree of root swelling or tubers | Swollen or fleshy | 1 | Water storage tissues to mitigate drought stress |
| | Absent | 0 | |
| **Chemistry** | | | |
| Photosynthetic pathway | CAM | 1 | Associated with improved photosynthetic performance under water stress and higher temperatures |
| | C4 | 0.5 | |
| | C3 | 0 | |

**Table S2.** Fog and drought functional trait adaptation scores for desert endemic study plants from the Central Desert.

| Trait | AA | HG | OH | PJ | PP | SH | SG |
|---|---|---|---|---|---|---|---|
| Maximum plant height | 0 | 0.5 | 0 | 0.5 | 1 | 0.5 | 0 |
| Leaf surface area | 0.5 | 0 | 1 | 0.5 | 1 | 0.5 | 0 |
| Leaf and stem hairs | 0.5 | 0.5 | 0 | 0 | 1 | 0.5 | 0.5 |
| Succulence | 0 | 0.5 | 0 | 0.5 | 0 | 0 | 0 |
| Rosettes | 0 | 0 | 0 | 0 | 0 | 0 | 0 |
| Photosynthetic stems | 0 | 0 | 0.5 | 0 | 0 | 0 | 0 |
| Phenology | 0.5 | 0 | 1 | 0 | 0 | 0.5 | 0 |
| Root swelling or tubers | 0 | 0 | 0 | 0.5 | 0 | 0 | 0 |
| Photosynthetic pathway[1] | 0.5 | - | - | - | - | - | - |
| Total | 1.5 | 1.5 | 2.5 | 2 | 3 | 2 | 0.5 |

[1]Insufficient information is available on the photosynthetic pathway of the study species, thus it is not included in the total score.

**Table S3.** Collated and retained environmental variables for environmental niche modelling of desert endemic plants.

| Variable | Description | Retained | Source |
|---|---|---|---|
| Elevation | Elevation | x | GTOPO30, US Geological survey 1996 |
| Aspect | Aspect of the focal raster cell | x | GTOPO30, US Geological survey 1996 |
| Slope | Slope of the focal raster cell | | GTOPO30, US Geological survey 1996 |
| Roughness | Difference between the maximum and the minimum value of a cell and its eight surrounding cells | x | GTOPO30, US Geological survey 1996 |
| TRI | Terrain Ruggedness Index: Mean of the absolute differences between the value of a cell and the value of its eight surrounding cells | | GTOPO30, US Geological survey 1996 |
| AnnualPET | Annual Potential Evapotranspiration: a measure of the ability of the atmosphere to remove water through evapotranspiration processes, given unlimited moisture | | Worldclim 2,Fick and Hijmans (2017); ENVIREM, Title and Bemmels (2018) |
| Aridity | Thornthwaite aridity index: index of the degree of water deficit below water need | x | Worldclim 2, Fick & Hijmans (2017); ENVIREM, Title and Bemmels (2018) |
| ClimMoistureIndex | A metric of relative wetness and aridity | | Worldclim 2, Fick & Hijmans (2017); ENVIREM, Title and Bemmels (2018) |
| Continentality | Average temp. of warmest month, minus average temp. of coldest month | | Worldclim 2, Fick & Hijmans (2017); ENVIREM, Title and Bemmels (2018) |
| EmbergerQ | Emberger's pluviothermic quotient: a metric that was designed to differentiate among Mediterranean type climates | | Worldclim 2, Fick & Hijmans (2017); ENVIREM, Title and Bemmels (2018) |
| MaxTempColdest | Maximum temperature of the coldest month | | Worldclim 2, Fick & Hijmans (2017); ENVIREM, Title and Bemmels (2018) |
| MinTempWarmest | Minimum temperature of the warmest month | | Worldclim 2, Fick & Hijmans (2017); ENVIREM, Title and Bemmels (2018) |
| PETColdQ | Mean monthly PET of coldest quarter | | Worldclim 2, Fick & Hijmans (2017); ENVIREM, Title and Bemmels (2018) |
| PETDriestQ | Mean monthly PET of driest quarter | | Worldclim 2, Fick & Hijmans (2017); ENVIREM, Title and Bemmels (2018) |
| PETseasonality | Monthly variability in potential evapotranspiration | | Worldclim 2, Fick & Hijmans (2017); ENVIREM, Title and Bemmels (2018) |
| PETWarmestQ | Mean monthly PET of warmest quarter | | Worldclim 2, Fick & Hijmans (2017); ENVIREM, Title and Bemmels (2018) |
| PETWettestQ | Mean monthly PET of wettest quarter | | Worldclim 2, Fick & Hijmans (2017); ENVIREM, Title and Bemmels (2018) |
| ThermicityIndex | Sum of mean annual temp., min. temp. of coldest month, max. temp. of the coldest month, *10. | | Worldclim 2, Fick & Hijmans (2017); ENVIREM, Title and Bemmels (2018) |
| BIO1 | Annual mean temperature (°C) | x | Worldclim 2, Fick & Hijmans (2017) |
| BIO2 | Mean diurnal temperature range (mean(period max-min)) (°C) | | Worldclim 2, Fick & Hijmans (2017) |
| BIO3 | Isothermality (Bio02 ÷ Bio07) | | Worldclim 2, Fick & Hijmans (2017) |
| BIO4 | Temperature seasonality (C of V) | | Worldclim 2, Fick & Hijmans (2017) |
| BIO5 | Max temperature of warmest week (°C) | | Worldclim 2, Fick & Hijmans (2017) |
| BIO6 | Min temperature of coldest week (°C) | | Worldclim 2, Fick & Hijmans (2017) |
| BIO7 | Temperature annual range (Bio05-Bio06) (°C) | | Worldclim 2, Fick & Hijmans (2017) |
| BIO8 | Mean temperature of wettest quarter (°C) | | Worldclim 2, Fick & Hijmans (2017) |

| | | | |
|---|---|---|---|
| BIO9 | Mean temperature of driest quarter (°C) | | Worldclim 2, Fick & Hijmans (2017) |
| BIO10 | Mean temperature of warmest quarter (°C) | | Worldclim 2, Fick & Hijmans (2017) |
| BIO11 | Mean temperature of coldest quarter (°C) | | Worldclim 2, Fick & Hijmans (2017) |
| BIO12 | Annual precipitation (mm) | | Worldclim 2, Fick & Hijmans (2017) |
| BIO13 | Precipitation of wettest week (mm) | | Worldclim 2, Fick & Hijmans (2017) |
| BIO14 | Precipitation of driest week (mm) | | Worldclim 2, Fick & Hijmans (2017) |
| BIO15 | Precipitation seasonality (C of V) | | Worldclim 2, Fick & Hijmans (2017) |
| BIO16 | Precipitation of wettest quarter (mm) | | Worldclim 2, Fick & Hijmans (2017) |
| BIO17 | Precipitation of driest quarter (mm) | | Worldclim 2, Fick & Hijmans (2017) |
| BIO18 | Precipitation of warmest quarter (mm) | | Worldclim 2, Fick & Hijmans (2017) |
| BIO19 | Precipitation of coldest quarter (mm) | | Worldclim 2, Fick & Hijmans (2017) |
| BIO20 | Annual mean radiation ($W\ m^{-2}$) | | Worldclim 2, Fick & Hijmans (2017) |
| BIO23 | Radiation seasonality (C of V) | | Worldclim 2, Fick & Hijmans (2017) |
| BIO28 | Annual mean moisture index | | Worldclim 2, Fick & Hijmans (2017) |
| BIO31 | Moisture index seasonality | | Worldclim 2, Fick & Hijmans (2017) |
| WS_mean | Mean annual wind speed | | Worldclim 2, Fick & Hijmans (2017) |
| WS_cv | Annual wind speed coefficient of variation | | Worldclim 2, Fick & Hijmans (2017); |
| MODCF_interannualSD | Within-year seasonality represented as the standard deviation of mean 2000-2014 monthly cloud frequencies | | Wilson and Jetz, (2016) |
| MODCF_intraannualSD | Mean between-year seasonality represented as the mean of the 2000-2014 monthly standard deviations | | Wilson and Jetz, (2016) |
| MODCF_meanannual | Mean annual cloud frequency (%) over 2000-2014 | x | Wilson and Jetz, (2016) |
| MODCF_seasonality | Timing of peak seasonal cloud concentration | x | Wilson and Jetz, (2016) |
| MODCF_spatialSD | Spatial variability of mean annual cloud cover represented as the standard deviation of mean annual cloud frequency within a one-degree moving window | | Wilson and Jetz, (2016) |
| Fog_mean | Mean brightness temperature difference for the period 2001-17 | x | MODIS, MCST (2018) |
| Fog_sd | Standard deviation of brightness temperature difference for the period 2001-17 | | MODIS, MCST (2018) |
| LST_mean | Mean night land surface temperature for the period 2001-17 (K) | x | MODIS, MCST (2018) |
| LST_sd | Standard deviation of night land surface temperature for the period 2001-17 (K) | | MODIS, MCST (2018) |
| Soil | Soil classification map | | FAO (2007) |

**Table S4.** Niche overlap of endemic Central Desert study species, estimated using Warren's I. Area is reported based on binary threshold occurrence.

| Species | AA | HG | OH | PP | SG | Area ($Km^2$) |
|---|---|---|---|---|---|---|
| *AA* | - | - | - | - | - | 2,383 |
| *HG* | 0.88 | - | - | - | - | 4,037 |
| *OH* | 0.91 | 0.90 | - | - | - | 8,530 |
| *PP* | 0.78 | 0.90 | 0.82 | - | - | 1,747 |
| *SG* | 0.95 | 0.87 | 0.87 | 0.71 | - | 565 |
| *SH* | 0.81 | 0.92 | 0.85 | 0.89 | 0.85 | 2,908 |

### Supplementary Figures

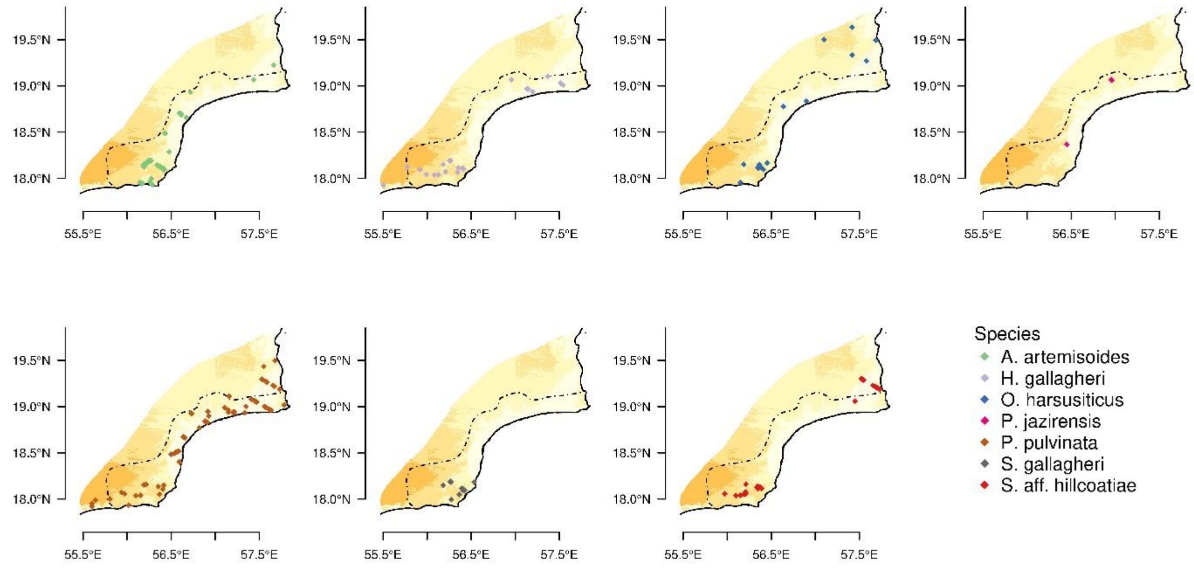


**Figure S1.** Species observation data for the southern Central Desert. Dashed line denotes focal study area.

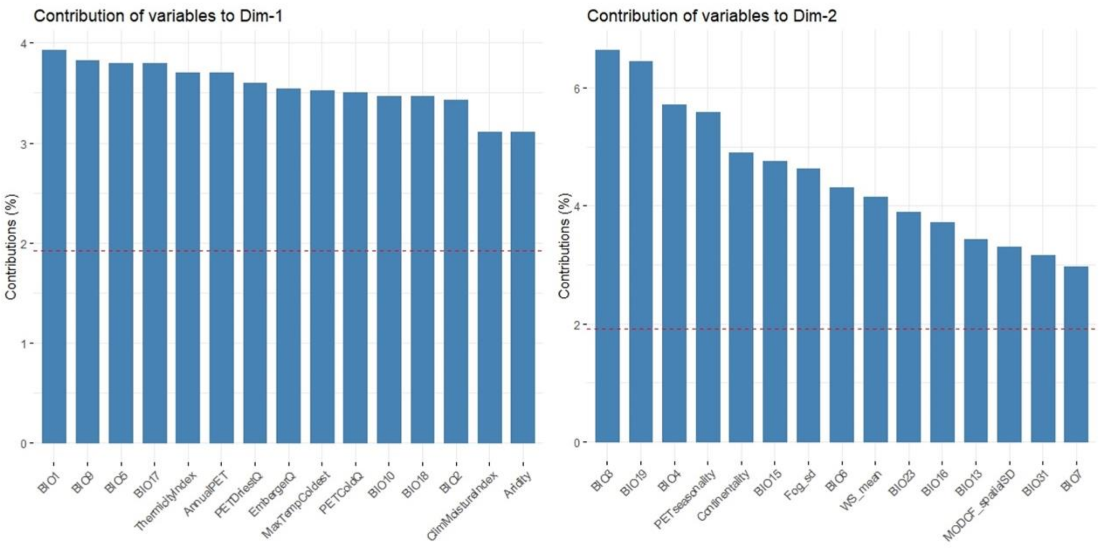


**Figure S2**. Top 15 variable loadings of the first and second principal component axes. Dashed line denotes expected value if all contributions were uniform.

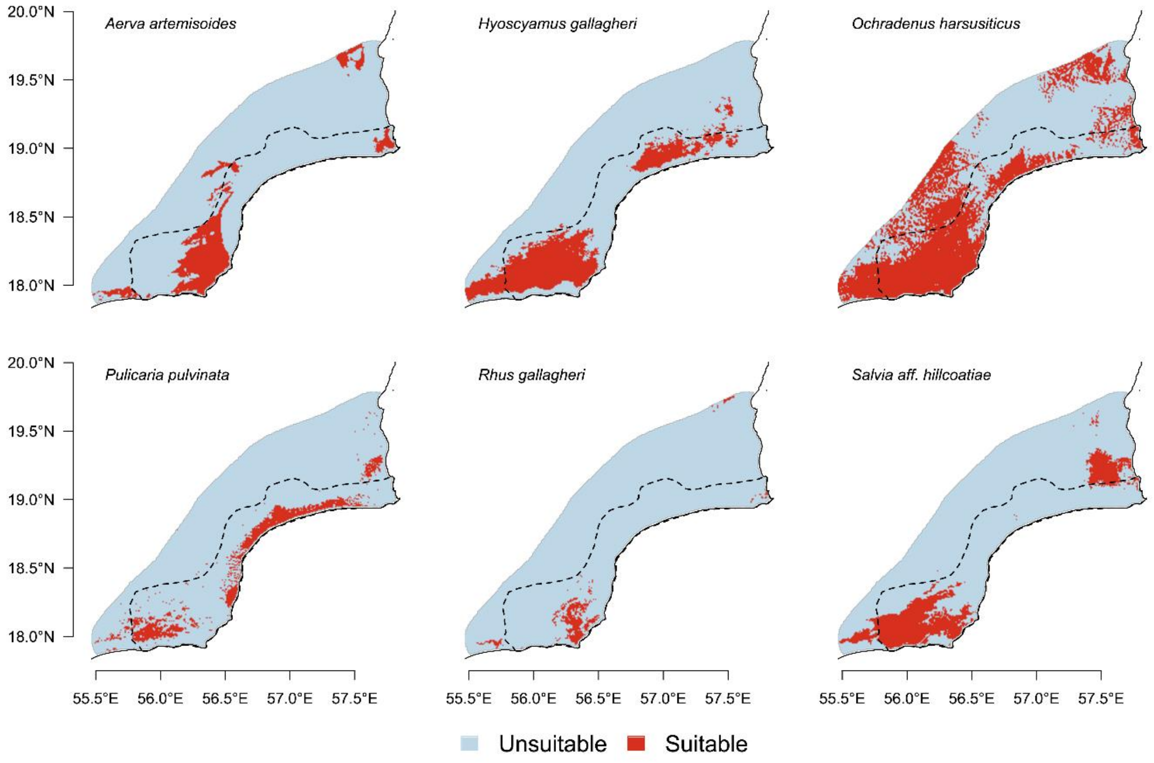


**Figure S3**. Binary threshold maps based on MESS and MTP values for six study species in the Central Desert.

#### References

Fick, S.E., Hijmans, R.J., 2017. Fick, Hijmans - 2017 - WorldClim 2 new 1-km spatial resolution climate surfaces for global land areas.pdf, International Journal of Climatology. https://doi.org/10.1002/joc.5086

Larraín-Barrios, B., Faúndez-Yancas, L., Búrquez, A., 2018. Plant functional trait structure in two fog deserts of America. Flora Morphol. Distrib. Funct. Ecol. Plants 243, 1–10. https://doi.org/10.1016/j.flora.2018.03.005

MODIS Characterization Support Team (MCST), 2018. MODIS 1km Calibrated Radiances Product.

Title, P.O., Bemmels, J.B., 2018. ENVIREM: an expanded set of bioclimatic and topographic variables increases flexibility and improves performance of ecological niche modeling. Ecography (Cop.). 41, 291–307. https://doi.org/10.1111/ecog.02880

Wilson, A.M., Jetz, W., 2016. Remotely Sensed High-Resolution Global Cloud Dynamics for Predicting Ecosystem and Biodiversity Distributions. PLoS Biol. 14, 1–20. https://doi.org/10.1371/journal.pbio.1002415
